# Dynamic membrane topology in an unassembled membrane protein

**DOI:** 10.1101/548537

**Authors:** Maximilian Seurig, Moira Ek, Gunnar von Heijne, Nir Fluman

## Abstract

Helical membrane proteins constitute roughly a quarter of all proteomes and perform diverse biological functions. To avoid aggregation, they undergo cotranslational membrane insertion and are typically assumed to attain stable transmembrane topologies immediately upon insertion. To what extent post-translational changes in topology are possible in-vivo and how they may affect biogenesis is incompletely understood. Here, we show that monomeric forms of Small Multidrug Resistance (SMR) proteins display topological dynamics, where the N-terminal transmembrane helix equilibrates between membrane-inserted and non-inserted states. We characterize the kinetics of the process and show how the composition of the helix regulates the topological dynamics. We further show that topological dynamics is a property of the unassembled monomeric protein, as the N-terminal helix becomes fixed in a transmembrane disposition upon dimerization. Membrane protein topology can thus remain dynamic long after cotranslational membrane insertion, and can be regulated by later assembly processes.

## Introduction

Helical membrane proteins constitute roughly a quarter of the proteome in all organisms ^1^ and are responsible for essential and diverse biological functions. Their prototypical primary sequence is characterized by stretches of hydrophobic sequence that form transmembrane helices (TMHs), separated by hydrophilic loops of varying lengths that protrude from both sides of the membrane. Their hydrophobic nature makes them aggregate quickly in aqueous solutions, necessitating their cotranslational insertion into the membrane. In this way, aggregation is avoided by sequential insertion of the individual TMHs as they are synthesized by the ribosome-translocon complex^2^. This cotranslational insertion sets the initial topology of the protein. Once inserted, it is typically assumed that the topology of a membrane protein cannot change anymore, mainly because any topological change would require the unfavorable transfer of hydrophilic loops through the membrane. Thus, membrane proteins are believed to follow a strict two-stage^2,3^ biogenesis mechanism: in the first stage, the protein cotranslationally inserts into the membrane and assumes its final, stable topology. In the second stage, the inserted TMHs pack together to form the final three-dimensional functional fold.

In recent years, however, several cases where topological dynamics does occur have been described. Some specialized proteins, such as bacteriophage holins and other pore-forming proteins, have evolved to switch topology when triggered by a physiological cue^4–6^. In other cases, topological dynamics was observed only under artificial conditions where, e.g., the lipid composition of the membrane was dramatically altered^7,8^, or the protein had to be truncated or mutated to observe dynamics ^9–13^. It remains unclear, however, if natural membrane proteins can display topological dynamics under physiological conditions and, if so, how such phenomena would affect the folding routes taken by membrane proteins *in vivo*. The biophysical principles underlying topological dynamics are also poorly understood.

Here we show that EmrE, a dimeric inner membrane protein from *Escherichia coli*, is topologically dynamic in its pre-assembled monomeric state *in vivo*. We find that, in monomeric EmrE, the N-terminal TMH1 flips in and out of the membrane, while dimeric EmrE has a stable membrane topology. We characterize the kinetics of the process and show how changes in the composition of TMH1 affect the rate of the topological dynamics. Our study suggests that membrane proteins may exist in a dynamic equilibrium between different topological states, and contributes to our understanding of how protein composition determines topology.

## Results

### The N-terminus of monomeric EmrE is partially mislocalizeds

EmrE is an *E. coli* homodimeric multidrug transporter, composed of two 4-TMH monomers^14^. It belongs to a unique group of proteins characterized by dual topology, where the monomer inserts into the membrane with two opposite orientations with equal probabilities^15–18^. The oppositely-oriented monomers then assemble to form the functional antiparallel dimer^16,17^. It has recently been shown that the dual topology of EmrE is generated early during biosynthesis when the monomers are inserted^18^, and that the overall orientations of monomers in the membrane are stable thereafter^19^.

Here, we studied the detailed topology of EmrE using a cysteine-accessibility assay. Briefly, the assay works by assessing the level of cysteine accessibility towards a periplasmic reagent (Fig. 1a)^19^. We engineered single Cys mutations in all loops and the two terminal tails of EmrE (Fig. 1b); all Cys mutants remained functional (Supplementary Fig. 1). We also tested the localization of the same loops/tails in well-characterized EmrE mutants engineered to insert with a single topology, either N_out_/C_out_ or N_in_/C_in_, in which both termini are located in the periplasm or cytoplasm, respectively (Fig. 1c, top). The N_out_/C_out_ and N_in_/C_in_ single-topology mutants cannot form the functional antiparallel dimer unless co-expressed^17,18,20^. To determine the locations of the individual cysteines relative to the membrane, cells were subjected to a 20 min reaction with AMS, a reagent that cannot cross the plasma membrane and therefore blocks only periplasmically accessible cysteines^19,21^ (Fig. 1a). The unblocked, cytosolic cysteines can later be reacted with maleimide-PEG, resulting in a shift in gel-migration. The results are quantified as fraction of AMS-blocked protein, reporting on the fraction of molecules that exposed the test Cys to the periplasm (see Supplementary Fig. 2).

**Fig. 1.**
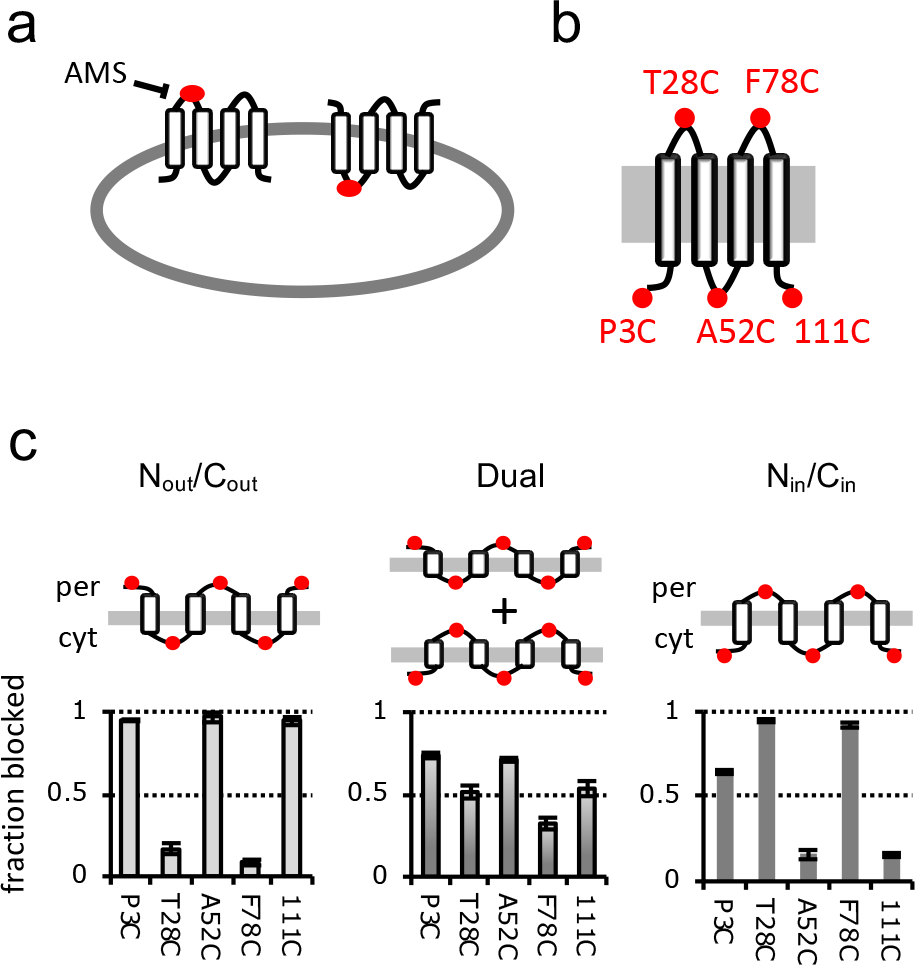
Topology scan of EmrE loops. **a**, Cysteine-accessibility assay. To determine the localization of an extramembrane loop (e.g. the loop connecting TM1 and TM2), a single Cys (red) is engineered into it. 4-acetamido-4’-maleimidylstilbene-2,2’-disulfonic acid (AMS), a sulfhydryl reagent that cannot cross the cytoplasmic membrane, is added to cells expressing the protein. The reagent only blocks cysteines in periplasmic loops, while cytosolic cysteines remain inaccessible. **b**, Locations of single-Cys substitutions in EmrE. Mutants used in the study contain only one Cys, to probe a unique loop every time. **c**, Periplasmic accessibility of the different loops as assessed by the fraction of EmrE molecules in which the Cys residue gets blocked by AMS. All loops were examined in the three topological backgrounds depicted on top. Error bars indicate standard deviations of independent experiments (n = 3).

For almost all positions, the results agree with the expected topology, namely the cysteines alternate between nearly full blocking or inaccessibility from the periplasm in the single-topology mutants, while showing partial blocking in the dual-topology wildtype protein (Fig. 1c). Unexpectedly, however, the N_in_/C_in_-P^3^C mutant showed ~65% periplasmic blocking, despite the expected cytosolic location of the N-terminal Cys (left bar in right-hand plot). We confirmed that the N_in_/C_in_-P^3^C mutant does not form intermolecular disulfides, which could have complicated the interpretation (Supplementary Fig. 3). It thus seems that a significant fraction of the N_in_/Cin molecules have the N-terminal tail mislocalized to the periplasm, indicating that TMH1 may be able to flip in and out of the membrane.

We next studied the dynamics of periplasmic exposure of the N-terminus in EmrE[N_in_/C_in_-P^3^C]. In order to follow the topology well after co-translational insertion has ended, a pulse-chase strategy was employed. Pulse-radiolabeled EmrE was chased for 5 min (with excess cold amino acids), ensuring that all radiolabeled EmrE molecules have finished synthesis several minutes before the topology assay. Wildtype EmrE is stably integrated in the membrane even after a much shorter time, without chase (Fig. 1c, and ref^19^). We then conducted kinetic blocking experiments by following the reaction of periplasmic AMS with the N-terminal Cys residue over time. In control experiments, we observed that the reaction of AMS with a Cys that is stably exposed to the periplasm is completed within 2.5 minutes (and probably much faster than this), while a cytosolic Cys remains mostly unblocked by AMS over the time-course of the experiment (Supplementary Fig. 4). The N-terminal Cys in EmrE[N_in_/C_in_-P^3^C] showed a remarkably different behavior, with 30% being blocked quickly and the remaining 70% gradually getting blocked over a 40-min period (Fig. 2a). By contrast, when confining the N-terminus to the cytosol by adding two N-terminal arginines^18^, we observed only ~5% blocking of the N-terminal Cys, with little increase over 40 min (Supplementary Fig. 5). We devised a simple kinetic model to explain the results (Fig. 2b), where the N-terminal TMH of EmrE(N_in_/C_in_) is assumed to reversibly equilibrate between a membrane-inserted and an uninserted state. In the uninserted state, the N-terminus is exposed to the periplasm, where it gets immediately and irreversibly blocked by AMS. To confirm that an equilibrium exists between inserted and uninserted TMH1, pulse-labelled EmrE[N_in_/C_in_-P^3^C] was incubated for 40 min prior to AMS blocking. The blocking kinetics was unaffected by the long pre-incubation, consistent with a pre-established equilibrium (Fig. 2c). The blocking kinetics was independent of AMS concentration (Fig. 2d), showing that the reaction with AMS is fast (*k*_*2*_ ≫ *k*_*−1*_) and that the overall blocking reaction is dominated by the rate-limiting transition from the inserted to the uninserted state (*k*_1_). Fitting the data to the simple kinetic scheme yielded a half-life of ~10 minutes for the transmembrane state (*k*_*1*_ = 1.1 ×10^−3^ ± 0.2×10^−3^ sec^−1^) and suggests that at equilibrium 31% (±3%) of the molecules have their N terminus (mis-) localized in the periplasm, enabling their immediate blocking upon AMS addition. We estimate a half-life of ~3 minutes for the reverse transition from the uninserted to the inserted state. TMH1 of monomeric EmrE is therefore not efficiently integrated into the membrane, but every molecule of EmrE samples the correct, fully inserted topology on a time scale of minutes.

**Fig. 2.**
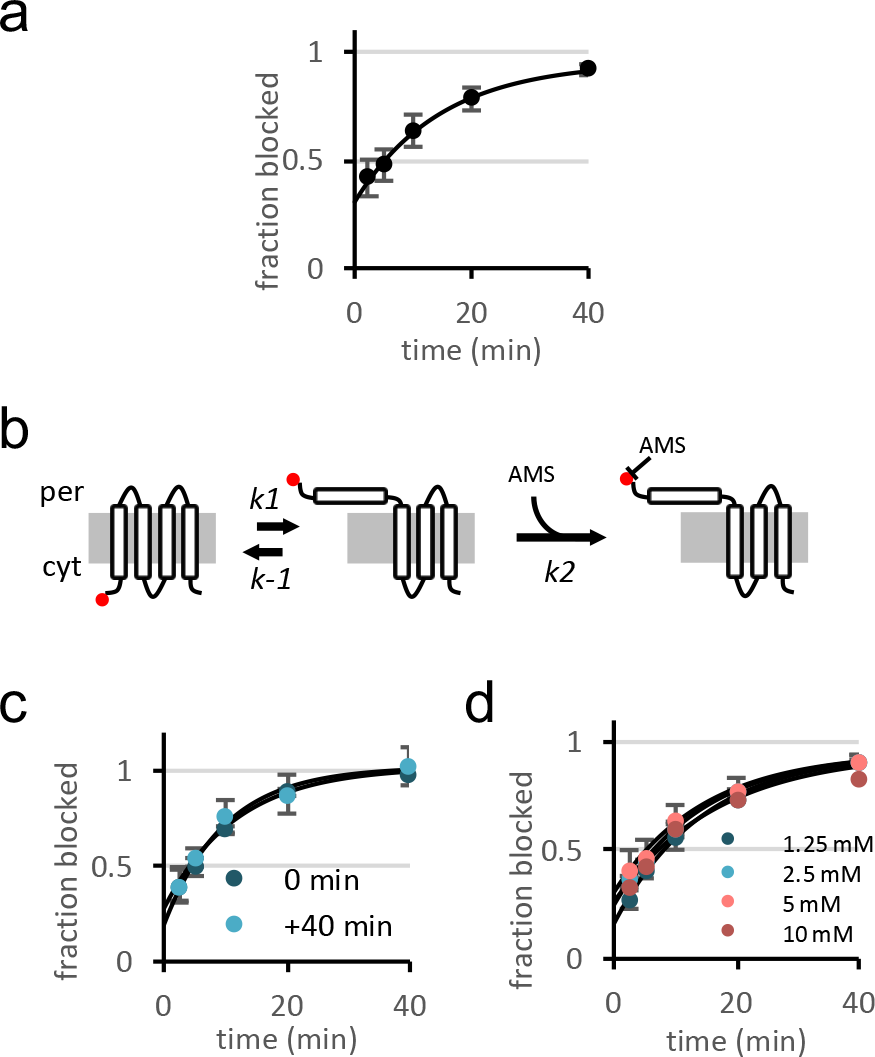
Topological dynamics of the EmrE N terminus. **a**, Periplasmic blocking of the N-terminus in EmrE[N_in_/C_in_-P^3^C] over time. The line is a fit to a single exponential kinetic equation (Y=Y_0_ + (Y_Plateau_−Y_0_)×(1−e^(−*k*x)^)). **b**, Kinetic model for periplasmic blocking of EmrE[N_in_/C_in_-P^3^C]. TMH1 equilibrates between an integrated, transmembrane state and a periplasmically accessible state. The latter is quickly and irreversibly blocked by periplasmic AMS. **c**, Effect of 40 min pre-incubation of EmrE[N_in_/C_in_-P^3^C] on the blocking kinetics, similar to (a). **d**, Effect of varying the AMS concentration on the blocking kinetics. Error bars indicate standard deviations of independent experiments (n ≥ 3).

### TMH1 is stabilized in the membrane upon dimerization

EmrE[N_in_/C_in_-P^3^C] does not have any oppositely oriented subunits to interact with when expressed alone. Therefore, the topologically dynamic species that we observe is monomeric EmrE. In the crystal structure of the EmrE dimer^16^, TMH1 makes extensive interactions with neighboring TMHs (Fig. 3a), making it hard to imagine that TMH1 could dynamically flip across the membrane. We therefore speculated that in the assembled dimer, the interactions with the TMHs would likely hold TMH1 in place, diminishing the topological dynamics. To test this, we reconstituted the functional dimer by coexpressing EmrE[N_in_/C_in_-P^3^C] with the oppositely oriented N_out_/C_out_ mutant. Indeed, dimerization dramatically slowed the dynamics, stabilizing the N-terminus in the cytosol, with the Cys protected from blocking over a long period of time (Fig. 3b). The slight blocking seen at long incubation times is similar to the background levels of negative controls (Supplementary Fig. 4b, Supplementary Fig. 5).

**Fig. 3.**
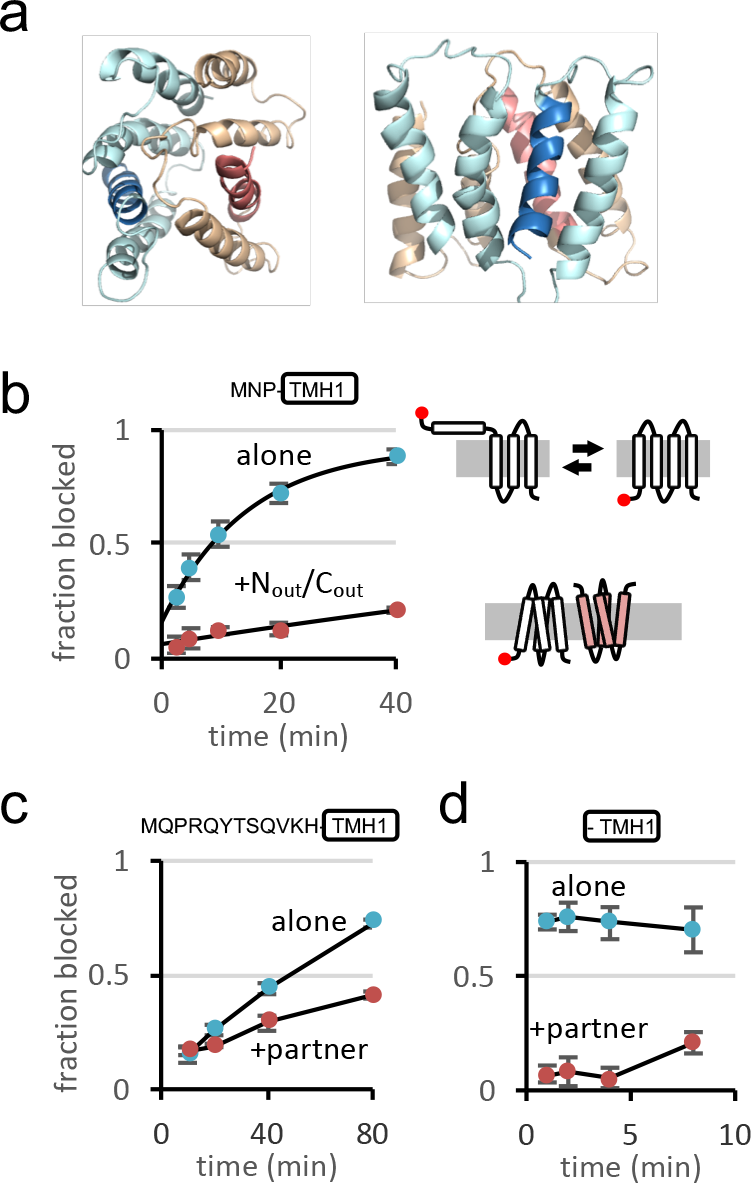
Effect of dimerization on topological dynamics. **a**, Structure of the EmrE dimer (PDB ID, 3B5D) viewed from above the membrane (left) or from within the membrane plane (right). The two protomers are colored light blue and orange, with TMH1 indicated by a darker color. **b**, Blocking kinetics of EmrE[N_in_/C_in_-P^3^C] when expressed alone, or coexpressed with the N_out_/C_out_ mutant. Topology diagrams depict the dynamic topological equilibrium of EmrE[N_in_/C_in_] (white) when expressed alone and the topological stability of the same protein when expressed with EmrE[N_out_/C_out_] (pink). The amino acid sequence preceding TMH1 of EmrE is shown above the graph. **c**, Blocking kinetics of an N-terminal Cys in the EmrE homolog Q1MPU8 when expressed alone or when coexpressed with its cognate SMR dimerization partner protein. The amino acid sequence preceding TMH1 is shown above the graph. **d**, Same as in (c), but for EmrE homolog B8J442. Note that B8J442 has no N-terminal tail sequence since TMH1 starts with the initiator Met. Error bars indicate standard deviations of independent experiments (n = 3).

We next asked if this phenomenon, of a monomer being topologically dynamic, is a rare curiosity, or if it could perhaps be more common than expected. To examine this, we turned to two distant homologs of EmrE (sharing only 15-30% sequence identity) that belong to the heterodimeric branch of the SMR family, B8J442 from *D. desulfuricans* and Q1MPU8 from *L. intracellularis* (names given are Uniprot accessions). Both proteins were shown before to have a N_in_/C_in_ topology and to dimerize with a N_out_/C_out_ partner of the SMR family that is encoded by a separate gene in their respective operons^19^. We introduced a reporter Cys in the cytosolic N terminus of both proteins and followed its periplasmic blocking by AMS over time. When co-expressed with their partner, both proteins showed a stable cytosolic location of the N terminus (Fig. 3 c,d). In contrast, when expressed alone, both proteins showed elevated periplasmic exposure. The periplasmic exposure of Q1MPU8 occurred quite slowly (30% over the dimer during 80 minutes Fig. 3 c), whereas that of B8J442 was apparently so rapid that it was predominantly blocked already at the 1 min time point (Fig. 3 d). Thus, monomers of the SMR family may frequently have a dynamic topology and dimerization is needed to achieve a stable, fully integrated state.

### Determinants of TMH1 insertion dynamics

What makes TMH1 dynamic while the other TMHs of EmrE remain fixed? A major barrier for topological dynamics is presumably the energetically costly transfer of hydrophilic extramembrane tail and loop regions across the membrane interior^22^. Indeed, the rate of N-terminal flipping in the three proteins that we have tested seems to correlate inversely with the length of the N-terminal hydrophilic tail. B8J442, which shows very fast periplasmic blocking, does not have a hydrophilic N-terminal tail at all, the exceedingly slowly blocked Q1MPU8 has a substantially longer hydrophilic tail, and EmrE has a short tail with only three residues, MNP, preceding the hydrophobic TMH1 (see sequences in Fig. 3b,c,d). To directly test the influence of the tail, we extended the N-terminus of EmrE[N_in_/C_in_-P^3^C] by one, two, or three serines. Indeed, the hydrophilic extensions gradually slowed the dynamics, nearly completely abolishing it already at two serines (Fig. 4a). Addition of serines also increased the fraction of membrane-inserted TMH1 at t = 0 (as indicated by the low level of blocking of the N terminus). This suggests that TMH1 is actually hydrophobic enough to get efficiently inserted into the inner membrane during the cotranslational insertion step (see later).

**Fig. 4.**
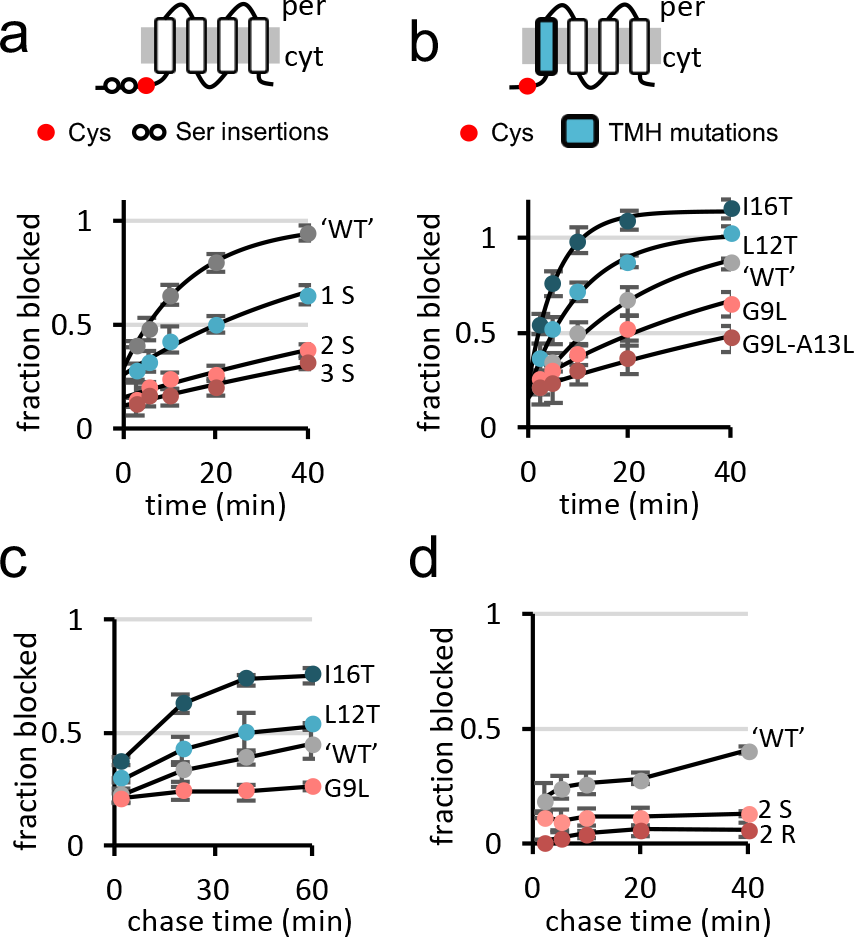
Effect of amino acid composition on TMH1 insertion dynamics. **a**, Effect of extending the N-terminal tail by 1-3 serine residues on the kinetics of EmrE[N_in_/C_in_-P^3^C] blocking by AMS. Curves are kinetic equation fits. **b**, Effect of mutations in TMH1 on the kinetics of EmrE[N_in_/C_in_-P^3^C] blocking by AMS. Hydrophilic and hydrophobic mutations are shown in blue and red, respectively. **c**, Effect of chase time on brief AMS blocking of EmrE[N_in_/C_in_-P^3^C] mutants with different TMH1 hydrophobicity. **d**, Effect of chase time on brief AMS blocking in mutants with 2 Ser (S) or 2 Arg (R) in the N-terminal tail of EmrE[N_in_/C_in_-P^3^C]. Error bars show ±SD of independent experiments (n ≥ 3).

We next wondered how the hydrophobicity of the membrane-embedded part of TMH1 may affect the dynamics. Making TMH1 more hydrophobic by one or two mutations gradually decreased the topological dynamics, while hydrophilic mutations had the opposite effects (Fig. 4b). Strikingly, ln(*k*_*1*_) was found to be directly proportional to the hydrophobicity of TMH1 as measured by the biological hydrophobicity scale^23^ (Supplementary Fig. 6). This suggests that the activation energy for the transition from the transmembrane to the periplasmically exposed state of TMH1 scales with its hydrophobicity. Collectively, it seems that hydrophilicity exerts opposite effects on the topological dynamics from within the TMH or outside of it: a hydrophilic loop slows topological dynamics, whereas a hydrophilic TMH accelerates it.

One puzzling observation was that even the most hydrophillic TMH1 mutants appear to be well inserted at t = 0, when the blocking reaction is initiated (Fig. 4b). This contradicted our expectation that hydrophilic TMH1 mutants will not be well inserted into the membrane. To follow the insertion status of TMH1 over time, we conducted a pulse-chase experiment. Cells were pulsed with radioactive [^35^S]-Met and then chased with nonradioactive Met, to allow the protein to age. At different aging times, we then briefly blocked periplasmic cysteines for two minutes only, in order to determine what fraction of the molecules had their N-terminus mis-localized to the periplasm. When the protein was still young (2 min chase), all mutants were roughly equally well inserted, with only 20% having their N-terminus accessible from the periplasm (Fig. 4c). Only over time did the hydrophobicity of TMH1 affect the insertion status of the different mutants, with hydrophilic mutants (I16T and L12T) equilibrating to the majority of the molecules having a periplasmically located N terminus, and the hydrophobic G9L mutant remaining well-inserted. Thus, membrane integration is favored during the initial cotranslational insertion step, even if the TMH is not thermodynamically stable in the membrane. This view is supported by a similar pulse-chase experiment with the mutant harboring two N-terminal serines. By halting topological dynamics, the hydrophilic tail extension caused TMH1 to remain stuck in its initial, well-inserted state (Fig. 4d), similar to a mutant where two arginines help fixing the N-terminus in the cytosol in accordance with the positive inside rule^24^. Collectively, these results suggest that TMH1 mutants that are not thermodynamically stable in the membrane nevertheless initially get efficiently integrated. A likely explanation is that the energetics of the initial cotranslational insertion step is dictated by an equilibration of TMH1 between the somewhat polar translocon channel and the surrounding membrane^23^. Once free of the translocon, TMH1 may then re-equilibrate between the transmembrane and periplasmically exposed state, unless a flanking hydrophilic tail traps it in the initial, membrane-integrated state.

## Discussion

Relatively little is known about how membrane proteins fold and mature *in vivo*. In the case of EmrE our studies suggest that the monomer is at least partially unfolded, such that TMH1 can move quite extensively relative to the rest of the protein. Stable folding appears to be coupled to dimerization and results in stabilization of TMH1 in its transmembrane disposition. We did not as yet find any functional role for the dynamics of the monomeric state, and its rate differs greatly between different homologs. *E. coli* EmrE likely dimerizes *in vivo* before appreciable re-equilibration of TMH1 has a chance to occur^25^, while for the B8J442 homolog, re-equilibration of TMH1 (Fig. 3d) and dimerization may possibly proceed on similar time scales. This suggests that the topological dynamics in this case has likely not been selected for by evolution, but probably was a coincidental byproduct of the need to conserve the functionally essential polar residue Glu^14^ in the middle of TMH1^13,14^. The dynamics was tolerated since the protein is still capable of populating the fully inserted state and form a stable functional dimer.

The existence of topological dynamics in monomeric EmrE demonstrates that membrane protein topology is not necessarily fixed after the cotranslational membrane-insertion step. This has implications for the folding and quality control routes taken by membrane proteins. For example, marginally hydrophobic transmembrane helices may have a chance to integrate post-translationally if they were missed during cotranslational insertion. Protein folding can perhaps even drive membrane insertion of such TMHs. A dynamic, mis-inserted state may be a hallmark of unfolding and perhaps also serve as a recognition determinant for cellular quality control^26^.

Membrane protein folding has traditionally been separated into two stages: TMH insertion and packing^3^. This in turn has simplified the description of the ‘second stage’ of folding, at which the already inserted helices pack together^27–30^. Current models assuming that the topology is completely fixed may need to accommodate more degrees of topological freedom of the unfolded state^22,31^. In that respect, better understanding of how common these phenomena are is crucial. Our analysis cannot yet answer this question but it can provide some hints. On the one hand, we observed topological dynamics in all three EmrE homologs tested, suggesting that it may not be extremely rare. On the other hand, out of the four TMHs of EmrE, only TMH1 is dynamic. The topological dynamics that we observe depends on the hydrophilicity of the TMH and its flanking tail. The short and not very polar N-terminal tail and the relatively weak hydrophobicity of EmrE TMH1 are not unusual among other TMHs in the *E. coli* inner-membrane proteome (Fig. 5a,b), but the combination of these two features makes EmrE TMH1 stand out (Fig. 5c). We propose that this combination is likely the major determinant of topological dynamics^22^. Interestingly, the analysis suggests that it is likely that several dozens, if not more, loops/tails in other membrane proteins are potentially dynamic. It is also possible that terminal TMHs may flip more easily than internal TMHs, since they are fixed to other parts of the protein only at one end. Overall, attaining the correct topology requires a balance between initial insertion, topological dynamics, protein folding, and oligomerization.

**Fig. 5.**
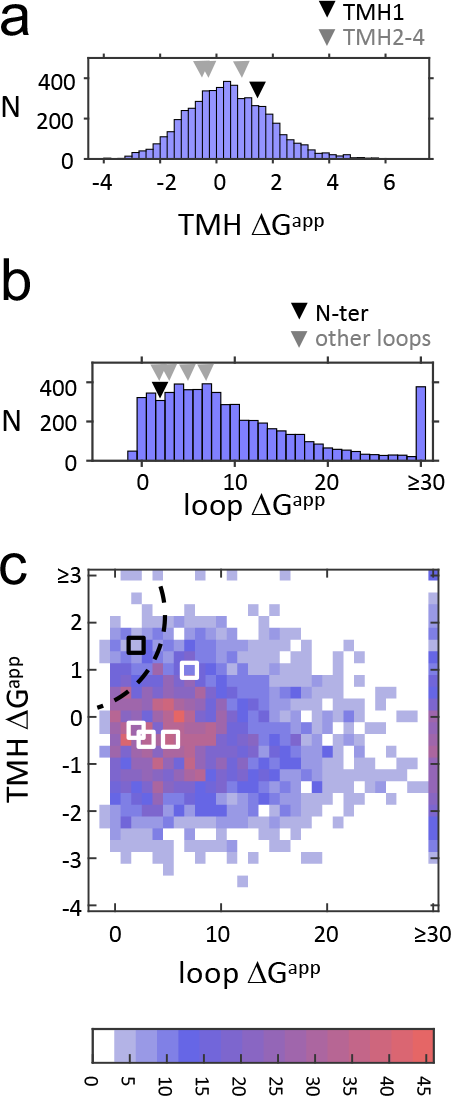
Hydrophobicity of TMHs and loops in the *E. coli* membrane proteome. ΔG^app^ values were calculated for all TMHs and extramembrane loops in 652 multispanning membrane proteins from *E. coli* (n = 5,286; a higher ΔG^app^ indicates a more hydrophilic sequence).

a. Distribution of TMH hydrophobicity among *E. coli* TMHs. Arrows show the respective values for TMH1 (black) and the other TMHs (gray) of EmrE.
b. Distribution of loop hydrophobicity among *E. coli* extramembrane loops. Arrows show the values for the N-terminal tail (black) and the other loops (gray) of EmrE.
c. Heatmap counting the paired occurrence of hydrophilicity of loops and their neighboring TMHs. Squares show the values for the N terminal tail (black) and the other loops (white) of EmrE. The dotted line indicates a region in the chart where TMH and loop hydrophilicity may potentially enable topological dynamics.

## Methods

All reagents were from Sigma unless otherwise noted.

### Bacterial strains

*E. coli* DH5α was used for plasmid propagation. For most protein expression experiments, *E. coli* BL21 (DE3) *ΔemrE::kana*^*R*^, *ΔmdtJI::Cm*^*R* 19^, deleted for the *emrE* gene and for the *mdtJI* operon was used to avoid potential interactions between expressed proteins and chromosomally-encoded SMR transporters. *E. coli* BL21 (DE3) *ΔdsbA::kana*^*R*^ was generated by P1 transduction of the resistance allele *ΔdsbA::kana*^*R*^ from the Keio collection^32^ into BL21(DE3) as described^33^.

### Plasmids

All proteins used were Cys-less versions of the original proteins. EmrE was expressed from pET-Duet-1 vector as described^19^, typically from the second MCS. When two EmrE variants were coexpressed, the cys-less N_out_/C_out_ mutant was expressed in excess from MCS1 which supports higher expression. The heterodimeric SMR transporters operons were expressed from pET19b (Novagen) as described^19^.

### Cell growth and specific radiolabeling of tag-less SMR proteins

Experiments were carried as described^19,34^. Briefly, BL21(DE3) *ΔemrE::kana*^*R*^ *ΔmdtJI::Cm*^*R*^ harboring plasmids encoding the proteins of interest were grown in M9 minimal medium at 37°C to mid-log phase (OD_600_ ~0.5). The cultures were then induced by 0.1 mM IPTG for 10 min, followed by 15 min incubation with 0.2 mg/ml rifampicin, during shaking at 37°C. Proteins were labeled with 15 μCi [^35^S]Met, typically for 5 min, then mixed with a high excess (2 mM) of non-radioactive methionine for variable chase times and put on ice for 5 min to stop the reaction. The chase was typically 5 min, except for Fig 1c, Supplementary Fig. 3 and Supplementary Fig. 2, where the chase time was 0 min (namely, the samples were put directly on ice after mixing with non-radioactive Met).

### Cysteine blocking in whole cells

The protocol was adapted from Fluman et al^19^. Radiolabeled cells (typically from a 10 ml culture for a kinetic profile) were washed by centrifugation at 4°C, 3,200 × *g*, 5 min and resuspension in 4 mL of Na-Mg buffer (150 mM NaCl, 30 mM Tris–HCl pH 7.5, 5 mM MgSO_4_, 1 mM TCEP (Tris(2-carboxyethyl)phosphine)). Cells were pelleted again and resuspended in 1 mL Na-Mg buffer and divided to three microcentrifuge tubes, containing 550, 110 or 220 μL, for blocking with AMS (Setareh Biotech), NEM, or water, respectively. Before each Cys blocking procedure, the respective culture was allowed to equilibrate to 30 °C for 4 min. At t=0, the sulfhydryl reagents were added (dissolved freshly, typically the AMS tube received 55 μL 100 mM AMS, the NEM tube received 22 μL of 0.2 M NEM, and the water tube 22 μL water). The cells were allowed to react for variable times at 30°C with gentle mixing, before quenching by mixing 100 μL cells with the pre-chilled tubes containing 0.5 mL Na-Mg buffer supplemented with 20 mM DTT. Cells were allowed to quench on ice for at least 20 min. Typically 5 timepoints with AMS were taken and one and two, respectively, early-range timepoints with NEM and water as controls. Samples were washed by centrifugation at 4°C, 10,000 × *g*, 2 min and resuspension in 0.9 ml of Na-Mg buffer. The cells were then pelleted again and resuspended in 150 μL of Lysozyme buffer (150 mM NaCl, 30 mM Tris–HCl pH 8, 10 mM EDTA, 1 mM TCEP, cOmplete protease inhibitor (Roche), 1 mg/ml lysozyme) and frozen in −20°C for at least 20 min. Cells were then lysed and the remaining unblocked cysteines were PEGylated.

### Cell lysis and cysteine PEGylation

Cells were disrupted by thawing at 25°C for 5 min followed by shaking at 37°C for 10 min. Then 0.9 ml of DNase solution (15 mM MgSO_4_, 10 μg/mL DNase I, 1 mM TCEP, 0.5 mM PMSF (Phenylmethanesulfonyl fluoride)) was added and the samples were allowed to shake at 37°C for 10 min before transferring them to ice. Crude membranes were collected by centrifugation at 4°C, 20,000 × *g*, 20 min and the pellet was resuspended in 50 μL of Na buffer (150 mM NaCl, 30 mM Tris–HCl pH 7.5, 1 mM TCEP). Membrane proteins were solubilized and PEGylated by adding 7 μL of 10% DDM (β-D-dodecyl maltoside, Anatrace) and 13 μL of 27mM mal-PEG (Methoxypolyethylene glycol maleimide, typically 5kDa from Sigma, or 2 kDa from Nanocs, NY, for protein Q1MPU8). The samples were incubated with continuous mixing at 30°C for 1 h and the reaction was quenched by mixing with 17.5 μL 5x Sample Buffer (120 mM Tris–HCl pH 6.8, 50% glycerol, 100 mM DTT, 2% SDS, and 0.1% bromophenol blue).

Samples were resolved on 15% SDS-PAGE gels run with a modified running buffer containing only 0.05% SDS ^35^. Gels were dried, visualized by autoradiography and quantified.

### Following N-terminal insertion status as the protein ages

BL21(DE3) *ΔemrE::kana*^R^ *ΔmdtJI::Cm*^R^ harboring plasmids encoding the proteins of interest were grown in M9 minimal medium at 37°C as above. Following the 15 min rifampicin treatment, a 7 mL culture was pulsed with 15 μCi [^35^S]Met and 1 mM TCEP for 1 min while shaking at 37°C. The culture was then chased by mixing with a high excess (2 mM) of non-radioactive methionine for variable chase times. At the end of the chase, cysteines were blocked by mixing 0.5 mL culture aliquots for 2 min at 37°C with 25 μL of 100 mM AMS, or 0.2 M NEM or water as controls. The reagents were then quenched by adding 25 μL of 1 M DTT, mixing for 30 sec in 37°C and then transferring to ice for at least 10 min. Samples were washed by centrifugation at 4°C, 10,000 × *g*, 2 min and resuspension in 0.9 ml of Na-Mg buffer. The cells were then pelleted again and resuspended in 150 μL of Lysozyme buffer and frozen in −20°C for at least 20 min. Cells were then lysed and the remaining unblocked cysteines were PEGylated as above.

### Data analysis

Gel bands were quantified using an in-house custom software and the fraction PEGylated and fraction of blocked cysteine were computed. Each data point of AMS-blocking was normalized to the NEM and water controls, corresponding to 100% and 0% blocking, respectively. When applicable, the data were fitted using nonlinear regression to the formula (Y(t)=Y_0_ + (Y_Plateau_−Y_0_)×(1−e^(−*k*t)^)). Where Y(t) is the fraction of blocked Cys at time t, Y_0_ is the fraction blocked upon an infinitesimally short reaction time with AMS, *k* is the rate constant (corresponding to *k*_*1*_ in Fig. 2b) and t is the time. Fitting was done using the Graphpad Prism software.

### Drug resistance assay

A single colony from a fresh transformation of BL21(DE3) *ΔemrE::kana*^R^, *ΔmdtJI::Cm*^R^, harboring the indicated pET plasmids was grown overnight in LB medium supplemented with 100 μg/ml ampicillin. The culture was then back-diluted 1:50 into the same medium and grown at 37°C to mid-logarithmic phase (OD_600_ ~0.5). Five 10-fold serial dilutions were then prepared from the cultures, with the highest density being of OD 0.1. The serial dilution was spotted (3.5 μL) on LB-agar-ampicillin plates containing 30 mM bis-tris-propane pH 7.0 and drug ethidium bromide (50 μg/mL). Identical plates without ethidium bromide served as control. The plates were allowed to grow for one or two nights at 37°C.

### Analysis of disulfides and their effect on cysteine blocking

Cultures (5 mL) of BL21(DE3) *ΔemrE::kana*^R^ *ΔmdtJI::Cm*^R^ or BL21(DE3) *ΔdsbA::kana*^R^ harboring a plasmids encoding EmrE[N_in_/C_in_-P^3^C] were grown and radiolabeled as described above, except that for the ‘plus TCEP’ condition cells received 1 mM TCEP together with the radiolabeled [^35^S]Met. After chilling the cultures on ice, 4 mL were used to measure Cys blocking in whole cells as described above (using 20 min blocking time). The remaining 1 mL was used to detect disulfide-bonded dimers as follows. Samples were washed once by centrifugation at 4°C, 3,200 × *g*, 5 min and resuspension in 1 mL of 150 mM NaCl, 30 mM Tris–HCl pH 7.5, 5 mM MgSO_4_. Cells were pelleted again and resuspended in 1 mL of the same buffer. NEM (100 μL, 0.2 M) was added and the samples were allowed to react at 30 °C with gentle mixing for 20 min. Samples were then washed twice by centrifugation at 4°C, 10,000 × *g*, 1 min and resuspension in 1 mL of 150 mM NaCl, 30 mM Tris–HCl pH 7.5, 5 mM MgSO_4_. The cells were then pelleted again and resuspended in 150 μL of Lysozyme buffer (150 mM NaCl, 30 mM Tris–HCl pH 8, 10 mM EDTA, cOmplete protease inhibitor (Roche), 1 mg/ml lysozyme, containing no TCEP) and frozen in − 20°C for at least 20 min. Cells were disrupted by thawing at 25°C for 5 min followed by shaking at 37°C for 10 min. Then 0.9 ml of DNase solution (15 mM MgSO_4_, 10 μg/mL DNase I, 0.5 mM PMSF, containing no TCEP) was added and the samples were allowed to shake at 37°C for 10 min before transferring them to ice. Crude membranes were collected by centrifugation at 4°C, 20,000 × *g*, 20 min and the pellet was resuspended in 50 μL of 150 mM NaCl, 30 mM Tris–HCl pH 7.5. Non-reducing 5x sample buffer (120 mM Tris–HCl pH 6.8, 50% glycerol, 2% SDS, and 0.1% bromophenol blue, 17.5 μL per sample) was added to all samples and they were then divided to two tubes, with or without 100 mM DTT. Samples were heated to 80 °C for 20 min and cooled down, then resolved on 15% SDS-PAGE gels run with a modified running buffer containing only 0.05% SDS ^35^. Gels were dried and visualized by autoradiography.

### Genome-wide analysis of TMH and loop hydrophilicity

The *E. coli* proteome was retrieved from UniprotKB/Swissprot database on November 2018. The transmembrane helix assignments were based on the database. Only proteins with 3 or more transmembrane helices were analyzed. The biological hydrophobicity^23^ was calculated for the entire protein sequences using TOPCONS^36^. In order to identify the most hydrophobic ΔG^app^ value for each TMH, the minimal ΔG^app^ value within ±5 residues from the TMH center was kept. In order to analyze loop hydrophilicity, loops sequences were extracted from the Uniprot entries and ΔG^app^ values for all of their residues were summed up. The ΔG^app^ values were taken from the biological hydrophobicity scale^23^, corresponding to residues positioned in the middle of the membrane. To calculate paired values of loop and TM hydrophilicity, the ΔG^app^ value of each loop was paired with the ΔG^app^ of a neighboring TMH. In cases where the loop has two neighboring TMHs, the lower TMH ΔG^app^ was taken.

## Acknowledgements

This work was supported by grants from the Knut and Alice Wallenberg Foundation (2012.0282), the Swedish Research Council (621-2014-3713), and the Swedish Cancer Foundation (15 0888) to GvH. NF was supported by long-term postdoctoral fellowships from EMBO/Marie Curie Actions (ALTF 211-2014) and from HFSP (LT000277/2015-L).

---

**Supplementary Fig. 1.**
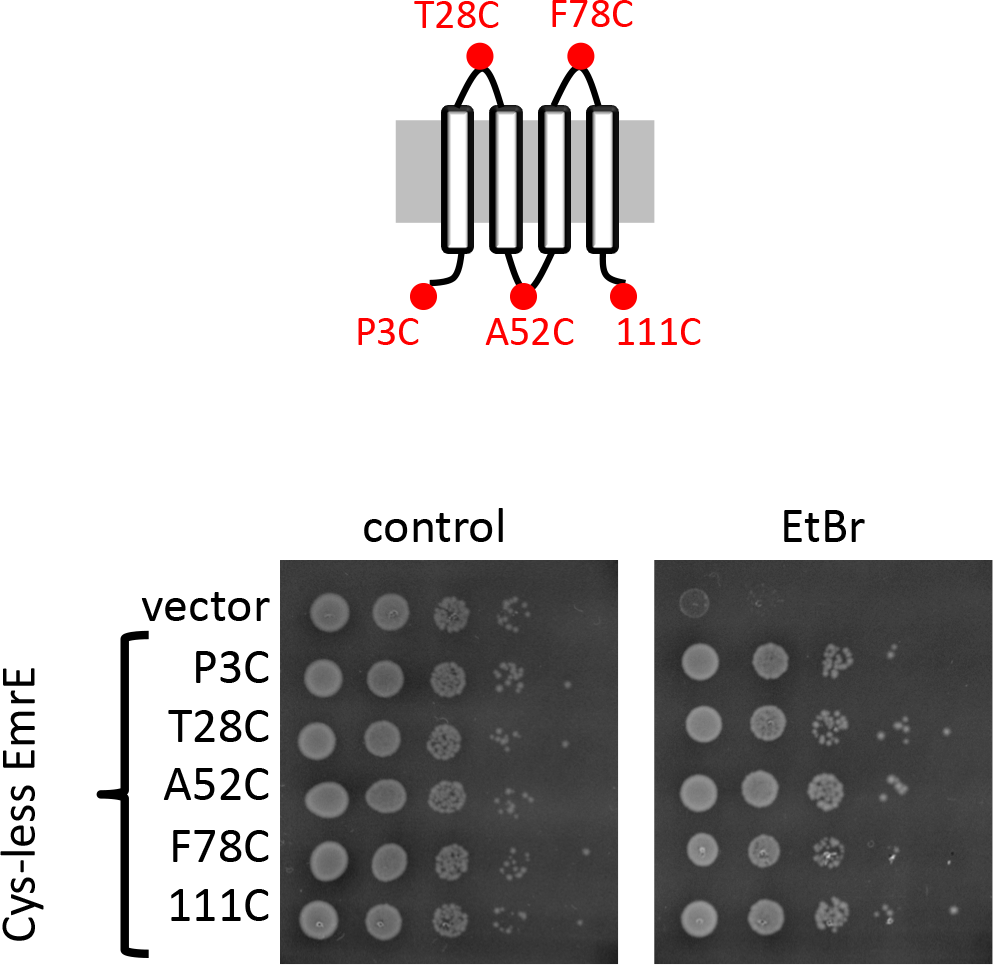
Ethidium bromide resistance conferred by EmrE single-Cys mutants. The efflux activity of EmrE mutants was inferred from their ability to support growth on toxic levels of the EmrE substrate ethidium bromide (EtBr). A serial 10-fold dilution of *E. coli* BL21(DE3) *ΔemrE::kana*^R^, *ΔmdtJI::Cm*^R^ harboring plasmids containing the indicated constructs was spotted on plates with or without EtBr (50 μg/mL). Cells harboring an empty vector serve as a negative control. Cells expressing all constructs grow well on control plates with no EtBr, whereas expression of functional EmrE is required to support growth on EtBr. Results are representative of 3 independent experiments.

**Supplementary Fig. 2.**
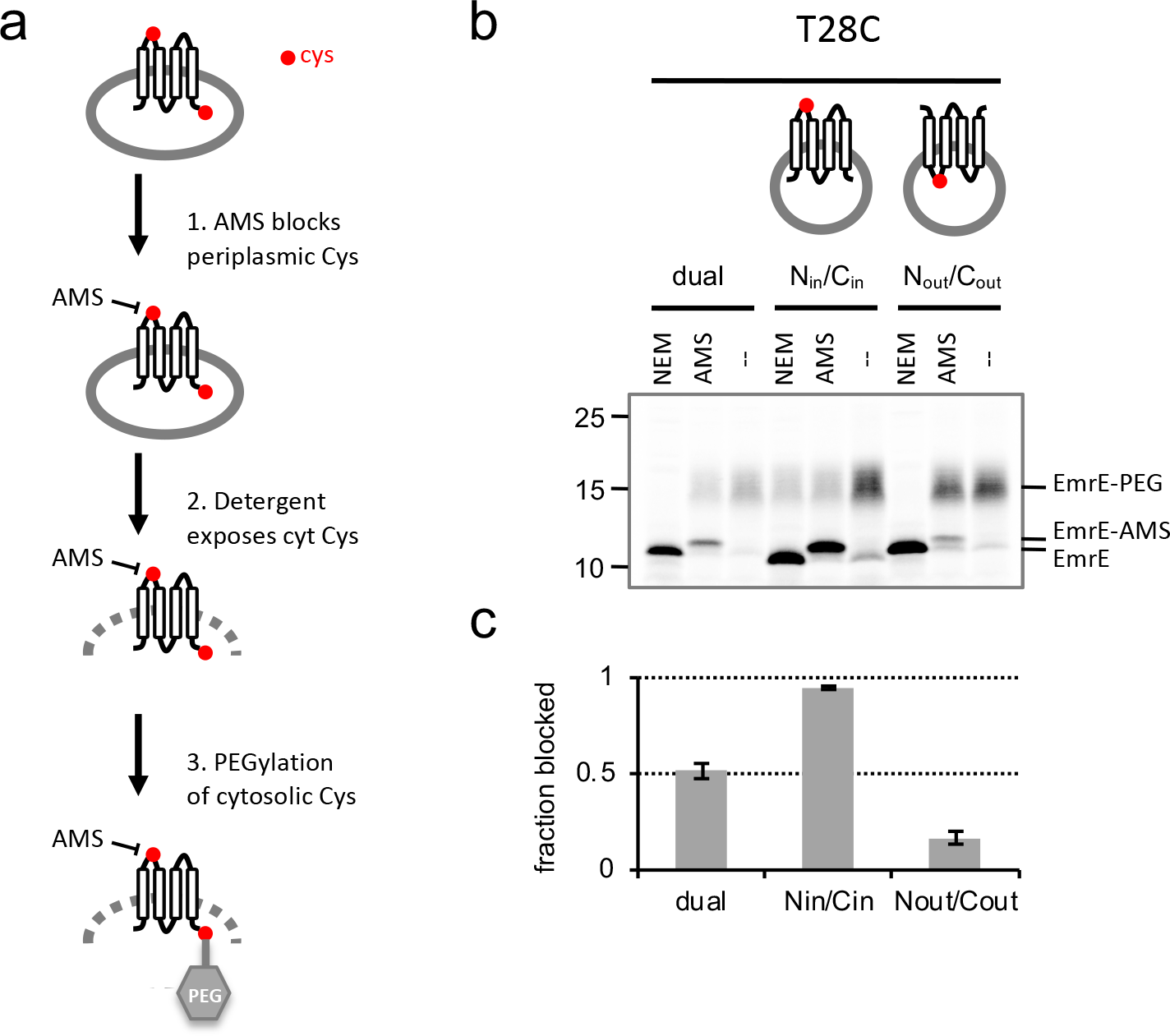
Topology assay using cysteine accessibility. a. Schematic description of the assay. Note that the EmrE mutants used in the assay typically contain a single Cys, unlike the two cysteines shown in the example. This depiction is shown here to demonstrate the difference between a periplasmic and cytosolic Cys. Brifely, a sulfhydryl reagent that cannot cross the cytoplasmic membrane, 4-acetamido-4ʹ-maleimidylstilbene-2,2ʹ-disulfonic acid (AMS), is added to whole cells to block any cysteines that are accessible from the periplasm. Subsequently, the membrane is disrupted and another reagent, maleimide-polyethylene glycol (mal-PEG), is added to derivatize any unblocked cysteines. Cytosolic cysteines remain reactive to mal-PEG, which increases the mass of the protein by 5 kDa. In contrast, periplasmic cysteines are blocked by AMS and cannot subsequently be PEGylated. As a control, a membrane-permeable reagent, *N*-ethylmaleimide (NEM), is used in the blocking step instead of AMS, to confirm that the test cysteines are in a solvent-exposed position that can be blocked from either the cytosol or the periplasm.
b. Analyzing the Cys accessibility of EmrE-T28C, in the dual topology protein or in mutants engineered to a single topology, either N_in_/C_in_ or N_out_/C_out_. The predicted localization of the Cys in position 28 relative to the inner membrane is shown above the gel. For each construct, lanes 1,2 and 3 show the amount of PEGylation remaining after blocking the Cys in whole cells using NEM, AMS and water (as control), respectively. NEM blocks all cysteines, whereas AMS blocks only periplasmic ones.
c. Densitometric quantification of the experiment in (b) to calculate the percent periplasmic blocking of Cys28 in the different topological mutants. For details on data analysis see Methods. The results confirm the expected locations of the Cys in the different constructs. Error bars indicate standard deviations from 3 independent experiments.

**Supplementary Fig. 3.**
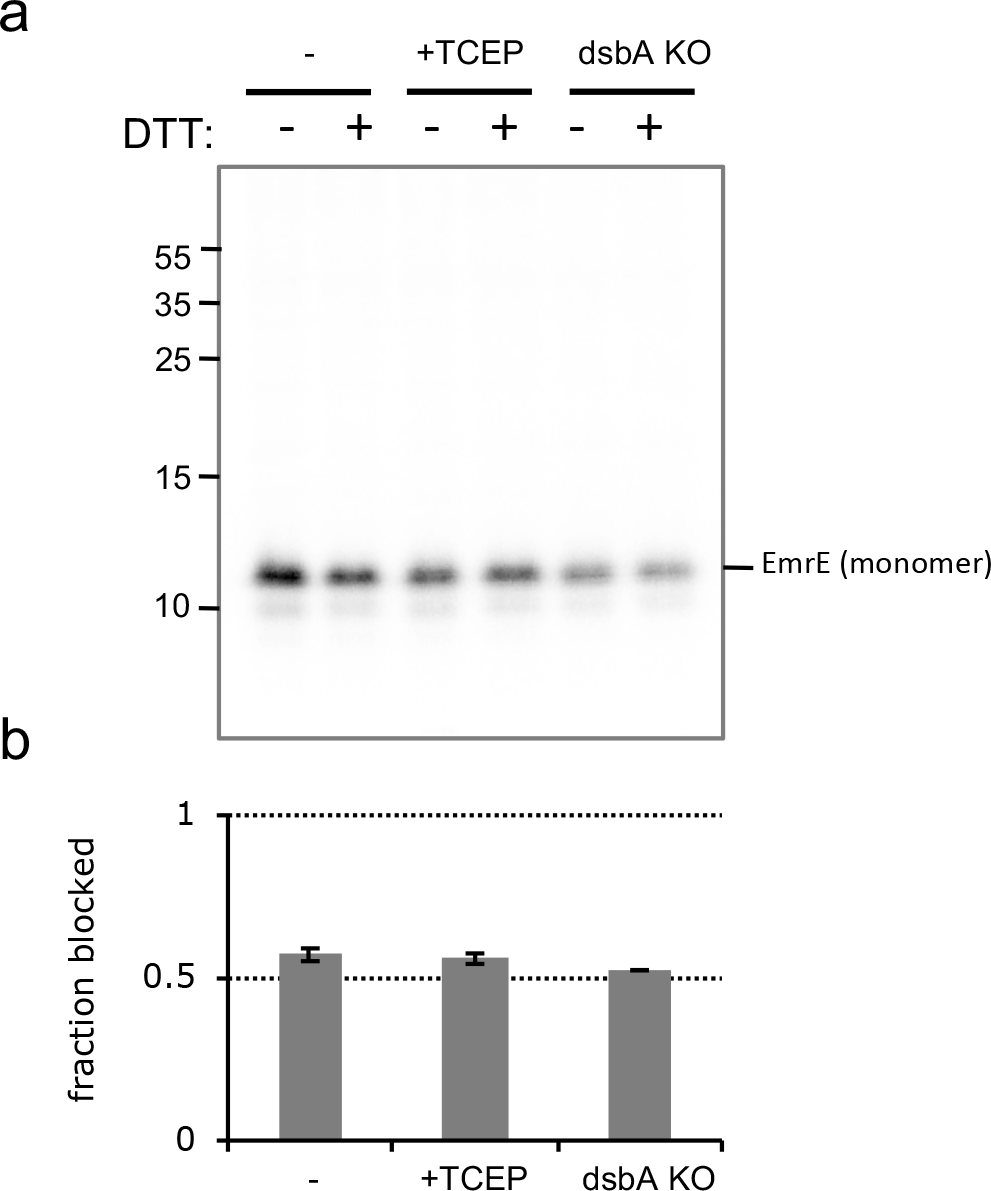
EmrE[N_in_/C_in_-P^3^C] does not form disulfides. EmrE[N_in_/C_in_-P^3^C] was expressed under three conditions: either identical to the topology-assay conditions (−) or in the presence of the reducing agent TCEP, or in an *E.* coli dsbA^−^ strain, in which the gene for the disulfide-catalyzing enzyme DsbA was knocked out.

a. No evidence for disulfide-bonded dimers of EmrE under any of the expression conditions. After expression, free cysteines were blocked by NEM in whole cells. Membranes were then isolated and boiled in sample buffer with or without the reducing agent dithiothreitol (DTT). SDS-PAGE analysis of the samples shows no evidence of disulfide-bonded EmrE.
b. The expression conditions do not impact the localization of the Cys in EmrE[N_in_/C_in_-P^3^C]. Cys-accessibility analysis of proteins expressed under the three conditions, indicates no difference in the level of periplasmic exposure of the test Cys. Error bars indicate standard deviation of independent experiments (n = 3).

**Supplementary Fig. 4.**
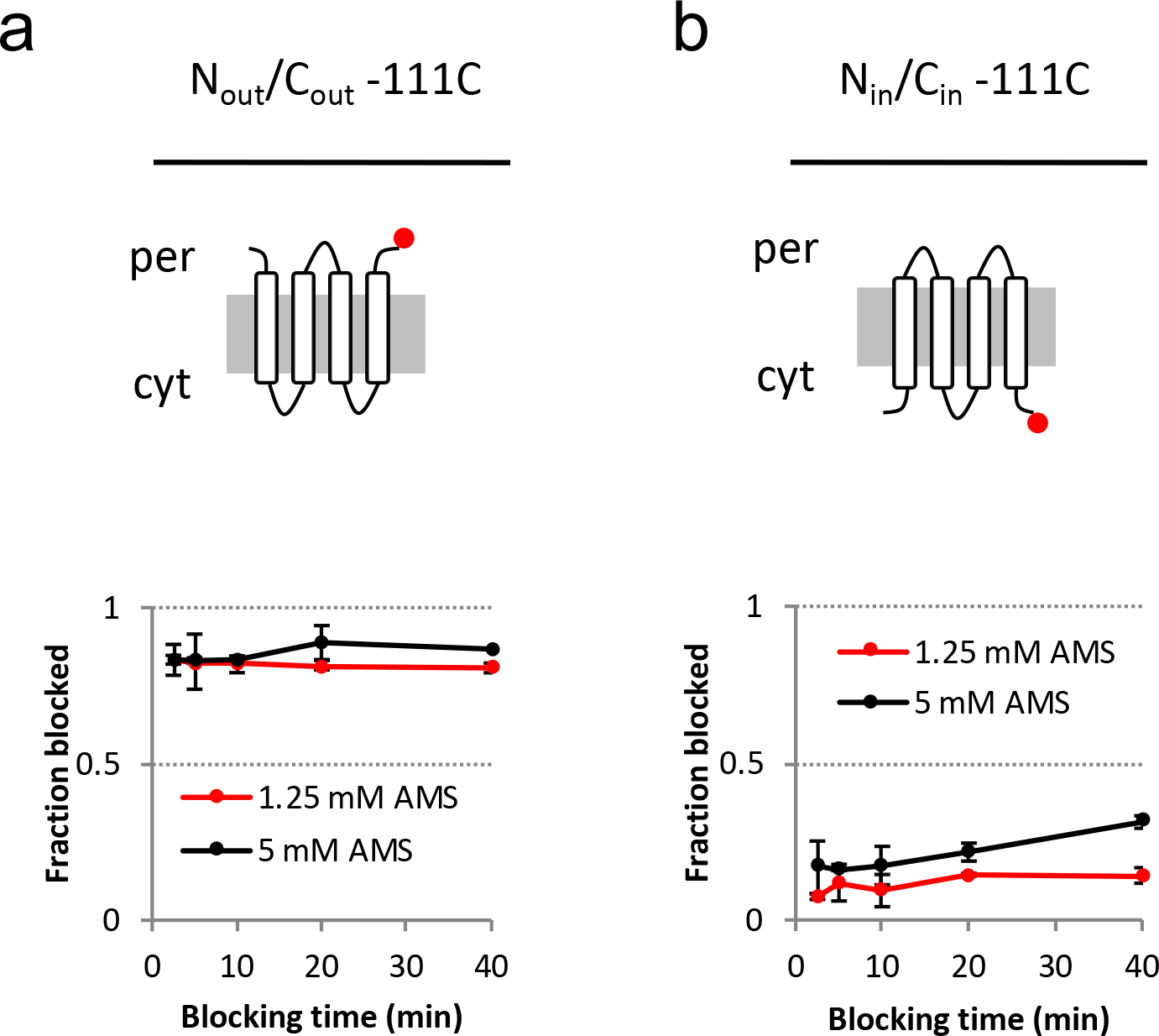
The C-terminal tail of EmrE does not have a dynamic toopology. Similar to Fig. 2a and d, the amount of periplasmic blocking of the C-terminus of EmrE over time was assessed, using EmrE mutants engineered to a single topology (N_out_/C_out_ or N_in_/C_in_) harboring a single Cys at the C-terminus (position 111). Two different AMS concentrations were used, similar to Fig. 2d. (a) The periplasmically exposed C-terminal Cys in EmrE[N_out_/C_out_-111C], gets quickly and nearly fully blocked. (b) The same Cys(at position 111), when confined to the cytosol in the N_in_/C_in_ mutant does not get appreciably blocked over time. Error bars indicate standard deviation of independent experiments (n = 3).

**Supplementary Fig. 5.**
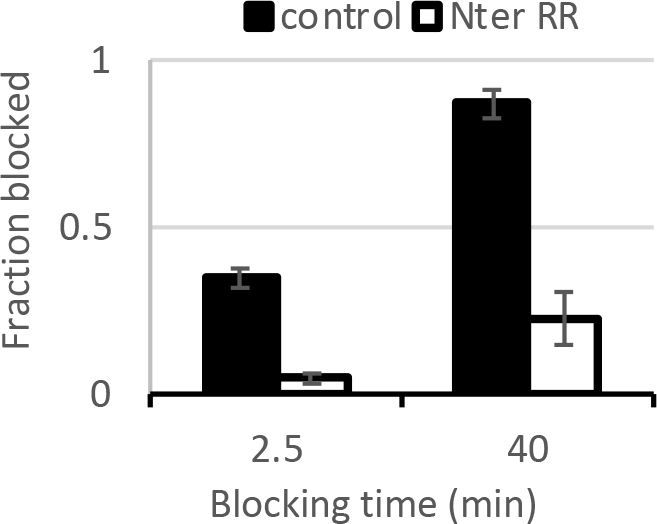
Addition of two N-terminal arginines locks the N-terminus of EmrE[N_in_/C_in_-P^3^C] in the cytosol. Periplasmic blocking of the N-terminus in EmrE[N_in_/C_in_-P^3^C](control) during 2.5 or 40 minutes blocking time. Nter RR is a similar construct containing two N-terminal arginines to hold the N terminus in the cytosol, in accordance with the positive-inside rule (see woodall et al. Ref 18). Error bars indicate standard deviation of independent experiments (n = 3).

**Supplementary Fig. 6.**
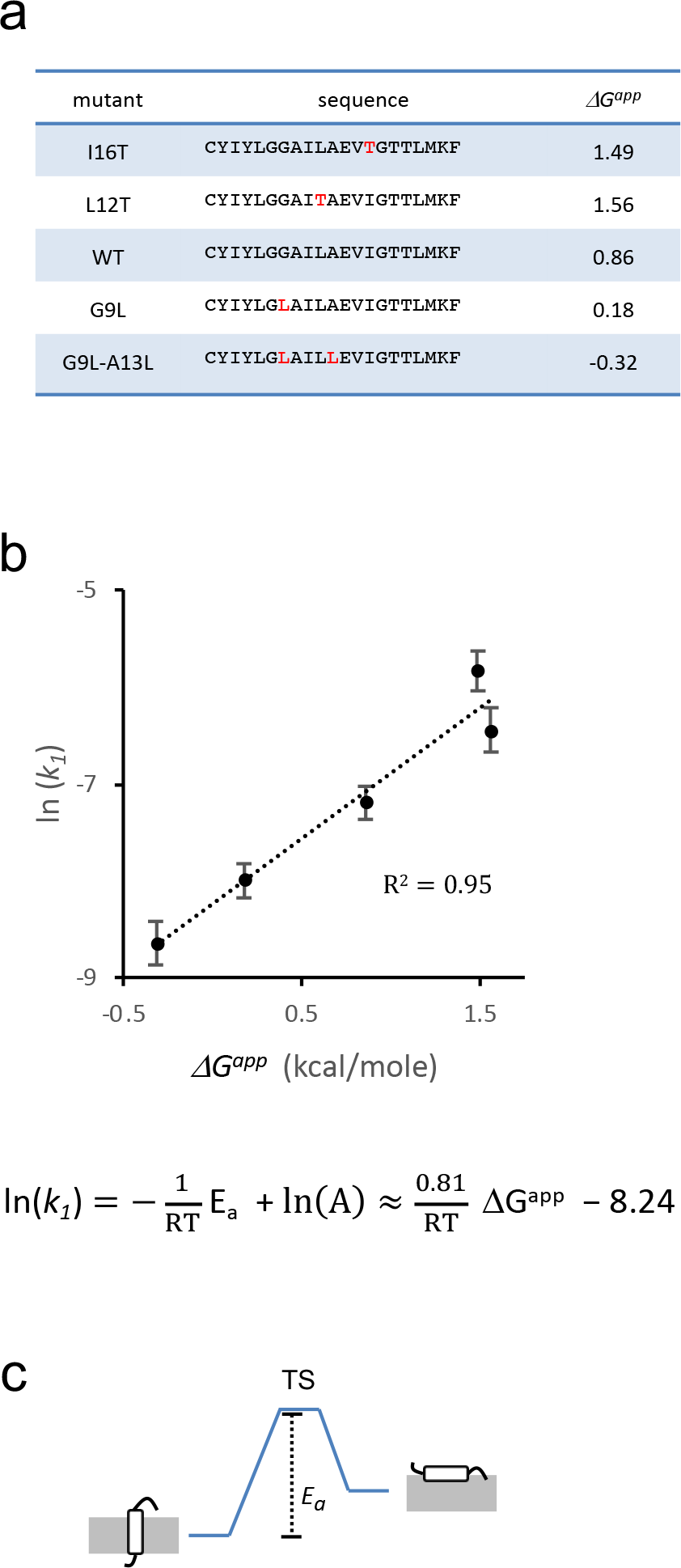
The Arrhenius activation energy for the transition from the transmembrane to the periplasmically exposed state of TMH1 scales with the hydrophobicity of TMH1. a. Hydrophobicity of TMH1 mutants (left), calculated by the *ΔG* predictor. The tool predicts the apparent *ΔG*^*app*^ for translocon-mediated membrane insertion (Hessa et al, 2007, ref 23).
b. The correlation between ln(*k*_*1*_) (see Fig. 2b) and Δ*G*^*app*^ implies that the Arrhenius activation energy *E*_*a*_ is proportional to *ΔG*^*app*^. The equation below is the Arrhenius equation, which indicates a linear relationship between the activation energy (*E_a_*) and ln(*k*) (*T,R* and *A* are temperature, the gas constant and the pre-exponential factor, respectively). Error bars indicate standard deviation of independent experiments (n = 3).
c. Energy diagrams for the activation energy (*E*_*a*_) for the transition from the transmembrane to the periplasmically exposed state of TMH1. TS, transition state.

